# Plant Virus Database: a resource for exploring the diversity of plant viruses and their interactions with hosts

**DOI:** 10.1101/2022.03.20.485054

**Authors:** Han Wu, Ping Fu, Qiong Fu, Zheng Zhang, Heping Zheng, Longfei Mao, Xiaoxu Li, Feng Yu, Yousong Peng

## Abstract

Plant viruses cause huge damage to commercial crops, yet the studies towards plant viruses are limited and the diversity of plant viruses are under-estimated yet. This study built an up-to-date atlas of plant viruses by computationally identifying viruses from the RNA-seq data in the One Thousand Plant Transcriptomes Initiative (1KP) and by integrating plant viruses from public databases, and further built the Plant Virus Database (PVD, freely available at http://47.90.94.155/PlantVirusBase/#/home) to store and organize these viruses. The PVD contained 3,206 virus species and 9,604 virus-plant host interactions which were more than twice that reported in previous plant virus databases. The plant viruses were observed to infect only a few plant hosts and vice versa. Analysis and comparison of the viromes in the Monocots and Eudicots, and those in the plants in tropical and temperate regions showed significant differences in the virome composition. Finally, several factors including the viral group (DNA and RNA viruses), enveloped or not, and the transmission mode of viruses, were found to have no or weak associations with the host range of plant viruses. Overall, the study not only provides a valuable resource for further studies of plant viruses, but also deepens our understanding towards the genetic diversity of plant viruses and the virus-host interactions.

## Introduction

Plant viruses are a type of DNA or RNA viruses that infect plants and often cause various types of diseases [1]. Although the diseases are rarely fatal to the plants, they can produce symptoms such as ringspot, mosaic pattern development, leaf yellowing and distortion, as well as deformed growth [2]. Currently, there is no effective cure for plant viral diseases [3]. The best way of controlling the viral disease is to destroy the infected plants, which usually results in huge damage to commercial crops [4].

The majority of plant viruses have the positive single-stranded RNA genome, with only a minority having other genome types [5,6]. The genome size of plant viruses is constrained as most plant viruses use plasmodesmata as means of entry into host cells [7,8]. The RNA plant viruses tend to have smaller genomes than DNA plant viruses [9]. Most plant virions are small due to the small genome size [6,8]. They come in two types: filaments and polygons [6]. Besides the viruses, the viroids and satellite viruses also infect plants and cause diseases [10]. The viroids consist of tiny single-stranded molecules of RNA, usually only a few hundred nucleotides long [11], while the satellite viruses are subviral pathogens that are entirely dependent upon their helper viruses’ replication machinery [12].

The interactions between viruses and plants are complex. As a consequence of long-term interactions, viruses and plants may co-evolve [13]. On the one hand, the plant virus can evolve through high mutation rates [8] for adaptation to plants which may cause diseases in plants; on the other hand, the plant could develop effective mechanisms to combat virus infection [14]. For example, the ribosome-inactivating proteins (RIPs), which are scattered all over the plant body, can inactivate viral proteins and play a key role in antiviral defenses [15]. The virus may change the hosts or increase the host range by evolution [16]. For example, the maize stripe virus has evolved to infect most maize genotypes during the past century [17]. The ecological factors and human activities may also increase the host range of plant viruses [16,18]. One of the most effective ways of establishing virus-host interactions is to introduce new host plant species and new viruses or their vectors into areas where they were not previously present before. For instance, shortly after the cacao was introduced from America, the Cacao swollen shoot virus emerged as an important pathogen of cacao in West Africa [18].

In recent years, novel viruses have been discovered at an unprecedented pace as a result of the rapid advancement of next-generation-sequencing (NGS) technology [19]. Numerous metagenomic and virome studies have been conducted to investigate the virome diversity in animals and humans. For example, Shi et al. used the RNA-seq technology to identify 1,445 RNA viruses from more than 220 invertebrates [20] and 214 distinctive and previously undescribed putative viral species from more than 186 vertebrates [21]. However, only a few virome studies have been conducted in the plants. For example, using the RNA-seq method, Lee et al. [22] identified a novel virus named Hevea brasiliensis virus during a virome analysis of Hevea brasiliensis. In another case, Rumbou et al. [23] used the RNA-seq approach to identify several novel viruses of the genera *Carlavirus, Idaeovirus* and *Capillovirus* from the birch trees of German and Finnish origin which exhibit symptoms of the birch leaf-roll disease. These findings highlight the importance of employing the RNA-seq method to identify new plant viruses. Numerous RNA-seq datasets from plants have been accumulated in the public databases. For example, the One Thousand Plant Transcriptomes Initiative (1KP) provided over 1,000 RNA-seq datasets from over 1,000 plant species [24]. These datasets can be used to identify plant viruses.

Several databases about plant viruses have been developed in previous studies, such as the DPVweb [25], Plant Viruses Online [26], EPPO-Q-bank Plant Viruses and Viroids database [27]. However, most of these databases are no longer maintained or updated, and contain limited viruses. To investigate the diversity of plant viruses, this study developed a computational workflow to identify plant viruses from the 1KP project; then, the plant viruses were manually curated from the NCBI GenBank database [28], the Virus-Host DB [29], and previous plant virus databases, and they were combined together and stored in the Plant Virus Database which was publicly available at http://47.90.94.155/PlantVirusBase/#/; finally, the evolution of viromes within the plants and the host range of plant viruses were analyzed. Overall, the study not only provides an up-to-date resource for plant viruses, but also deepens our understanding towards the genetic diversity of plant viruses and the virus-host interactions.

## Materials and methods

### Identification of viruses from the 1KP project

A total of 1,344 RNA-seq datasets, which cover 1,173 green plants and chloroplast-bearing species, were downloaded from the 1KP project [24]. A computational workflow was developed to identify viruses from the RNA-seq datasets (Figure 1). Firstly, the tool of fastp (version 0.21.0) [30] was used to trim low-quality reads. Then, the remaining reads were assembled into contigs using the Trinity program (version 2.1.1) [31] with default parameter settings. The assembled contigs were quantified using the RSEM software (version 1.3.0) [32]. Then, the contigs were queried against a library of plant virus protein sequences retrieved from the NCBI RefSeq database [33] on Jun 10, 2020 using BLASTX (version 2.10.1+) [34]. The contigs with an E-value of less than 1E-5 to the best hit were labeled as hypothetical plant viral contigs. The taxonomy of the best hit was assigned to the queried contig. For other contigs, they were further queried against the protein sequence of the RdRP domain (cd01699) derived from the Conserved Domain Database (version 3.18) [35] with RPSBLAST (version 2.10.1+). Contigs with one or more TPM, an E-value of less than 1E-2, a sequence identity and a coverage rate of greater than 0.3, and an alignment length of more than 100 to the best hit, were identified as hypothetical novel plant viral contigs. Finally, the hypothetical plant viral contigs were queried against the NR database [36] (downloaded on November 17^th^, 2020) with BLASTX to remove the false positives. The viral contigs with the best hit belonging to plant viruses were kept and considered as plant viral contigs.

**Figure 1.**
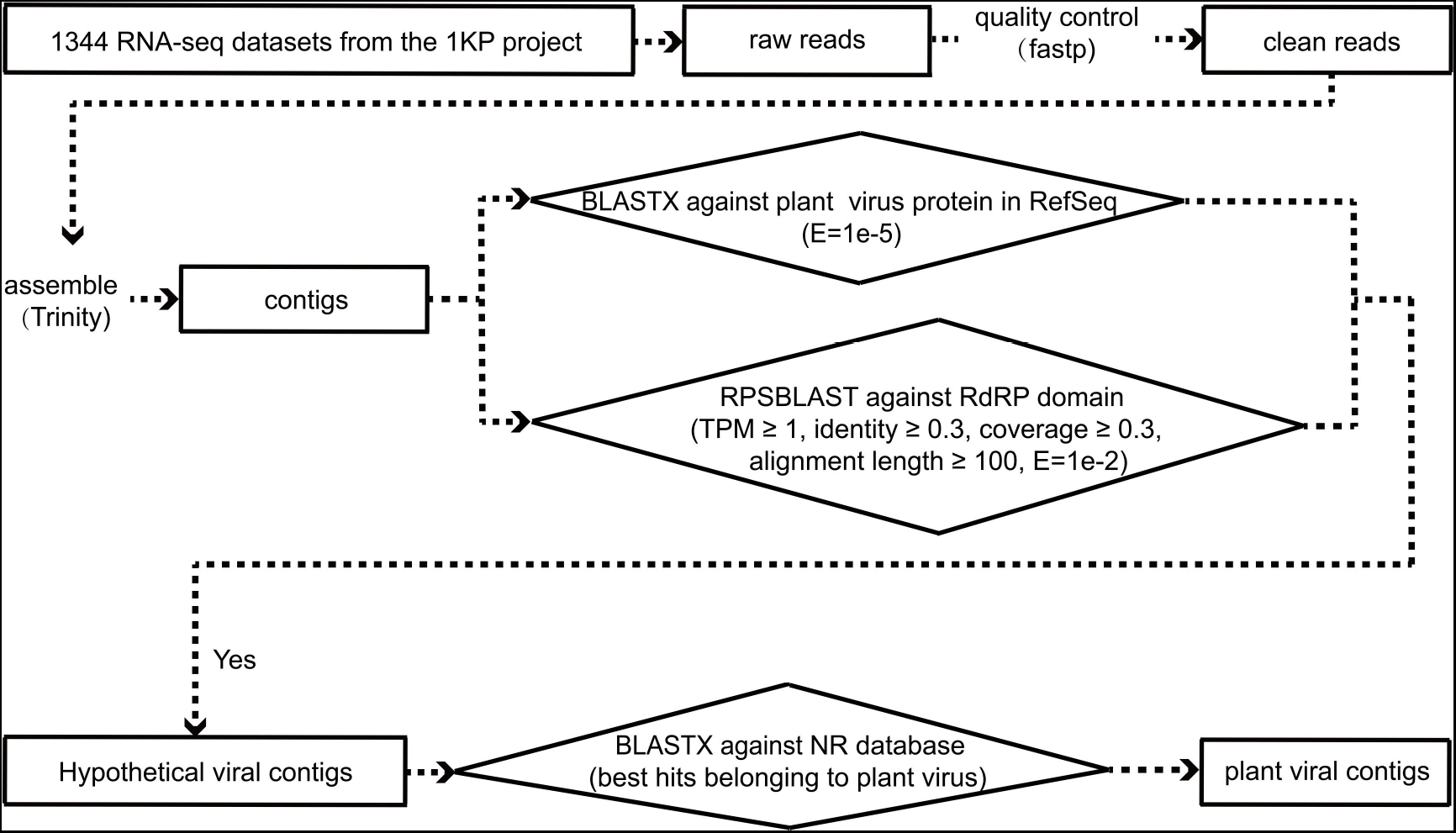
The workflow of identifying viruses from the 1KP project.

### Integration of viruses from public databases

In addition to the virus identified from the 1KP project, the plant viruses were also curated from the following sources: 1) the NCBI GenBank database. All viral sequences were firstly downloaded with the GenBank format from the NCBI GenBank database on September 14^th^, 2020. Viruses that were isolated from plant samples were obtained with the help of Taxonkit [37]. Viruses that have plants registered as their hosts were defined as plant viruses. Only the plant viruses were kept for further analysis. The tissue from which viruses were isolated was obtained from the field of “isolation_source”; 2) the Virus-Host DB. The Virus-Host DB is a database that provides data about the relationships between viruses and their hosts. The virus-plant interactions were directly downloaded from the database; 3) DPVweb. The virus-plant interactions were also directly downloaded from the database. The virus-plant interactions obtained above were merged and de-duplicated at the species level.

### Quantify the host range of viruses

To quantify the host range of viruses, firstly, a common tree was built for all plant hosts of viruses by the NCBI Taxonomy database [38]; then, the length of each branch in the common tree was changed to 1, and the distances between any pair of nodes were calculated with a Perl script. For viruses that infected plant hosts within a genus, the host range of them was calculated by dividing the number of plant hosts by the number of plants in the genus; for other viruses, the host range was calculated as the maximum distance of all pairs of plant hosts.

### The geography distribution of plant virus hosts

We focused on the angiosperm since it accounted for more than 95% of virus hosts. The geographical distribution of each plant family was manually curated from Wu et al. [39] and the Angiosperm Phylogeny Group’s studies [40].

### Technical details of the Plant Virus Database

The front-end was coded in standard HTML/JavaScript/CSS and deployed on the nginx (version 1.6.2), which is a high-performance HTTP server and reverse proxy. The back-end was coded in Python (version 3.6) that runs on the tornado web server (version 5.1), which is a Python web framework and asynchronous networking library. The JavaScript Object Notation (JSON) format was employed to implement the communication between the client-side and server-side layers. The data were stored in a MySQL (version 5.5.62) database.

### Statistical analysis

All the statistical analyses were conducted in R (version 3.2.5). The Wilcoxon rank-sum test was conducted by the function of *wilcox.test()* in R. The correlation coefficient was calculated by the function of *cor.test()* in R.

### Data availability

All the virus-plant interactions used in the study were publicly available at the Plant Virus Database which is freely available at http://47.90.94.155/PlantVirusBase/#/home.

## Results

### Virus identification from the 1KP project

The 1KP project (NCBI Accession Number: PRJEB4922) included a total of 1,344 RNA-seq datasets which summed up to 1.9 TB. A computational workflow (Figure 1, see Materials and Methods) was developed to identify viral sequences from the RNA-seq data in the 1KP project. A total of 270 virus species from 29 viral families were identified in 335 plant species from 76 orders. However, no viruses were identified in 73% of RNA-seq samples. No more than five virus species were detected in most of the remaining RNA-seq samples (Figure S1).

The viral families of *Potyviridae, Caulimoviridae, Geminiviridae, Partitiviridae, Secoviridae* and *Alphaflexiviridae* were most frequently observed, accounting for 61% of all virus species identified (Figure S2A). 61% of virus species were only identified in one plant species, and 96% of virus species were identified in less than ten plant species (Figure S2B). The orders of Lamiales, Caryophyllales, Malpighiales and Asparagales were found to have most plants (Figure S2C). An average of only 1.9 virus species were identified from each plant species (Figure S2D). Furthermore, no novel plant viruses were discovered in the 1KP project.

### Integration of plant viruses from public databases

To investigate the diversity of plant viruses, besides the plant viruses identified in the 1KP project, the plant viruses in public databases including NCBI GenBank, Virus-Host DB and DPVweb were integrated together (see Materials and Methods). A total of 9,604 pairs of virus-plant interactions were obtained, including 3,206 virus species from 46 viral families and 2,231 plant species from 92 orders. Most of these interactions (83%) were obtained from the NCBI GenBank database (Figure 2); 3,706 and 1,276 pairs of interactions were obtained from the DPVweb database and the Virus-Host DB, respectively; only 629 pairs of virus-plant interactions were obtained from the 1KP project. Notably, most of virus-plant interactions identified in the 1KP project were not previously reported in other databases.

**Figure 2.**
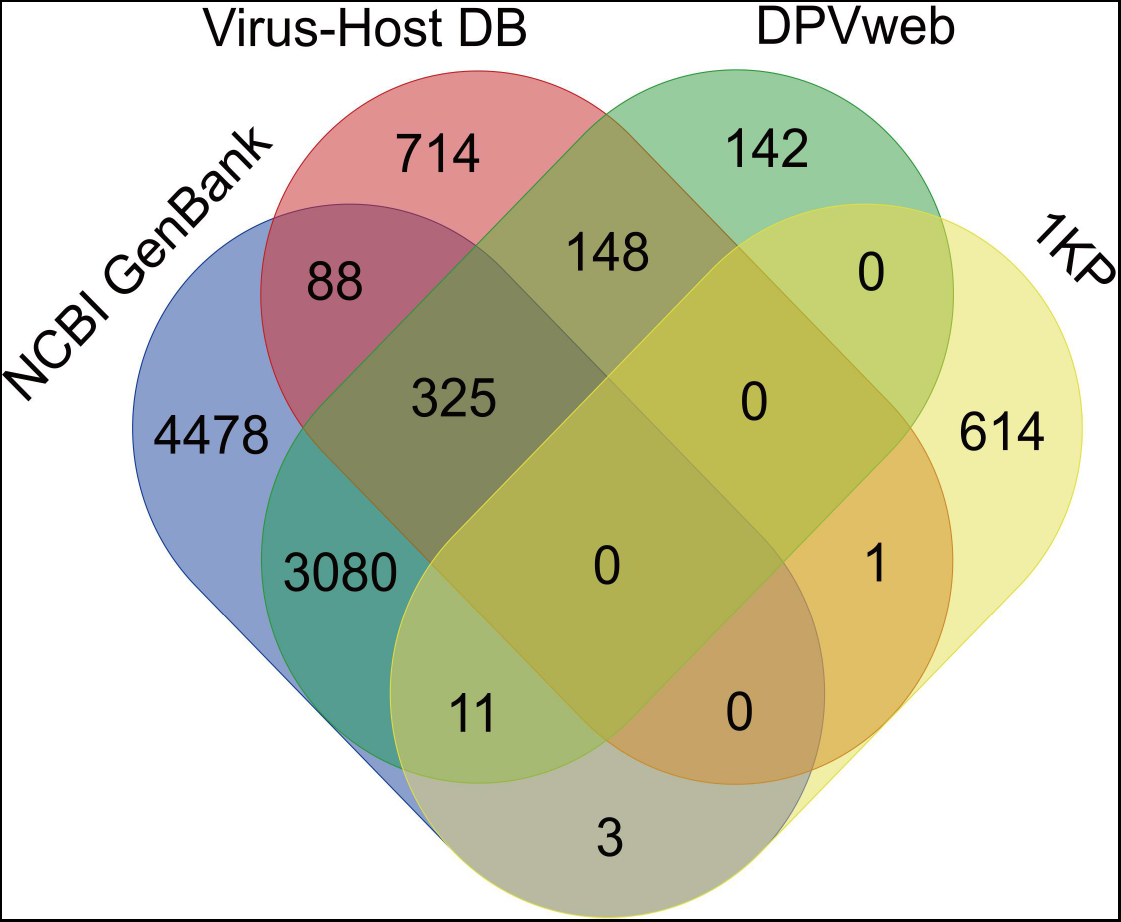
Venn diagram of sources of the virus-host interactions used in this study.

The 3,206 viral species belonged to 46 viral families, among which the *Geminiviridae* contained the largest number of viral species. The top five viral families contained 1,579 viral species which accounted for about half of all virus species (Figure 3A). Over 2,000 virus species only infected one plant species and 94% of virus species infected less than 10 plant species (Figure 3B). Some virus species infected more than 10 plant species. For example, the Cucumber mosaic virus infected 276 plant species across 31 orders.

**Figure 3.**
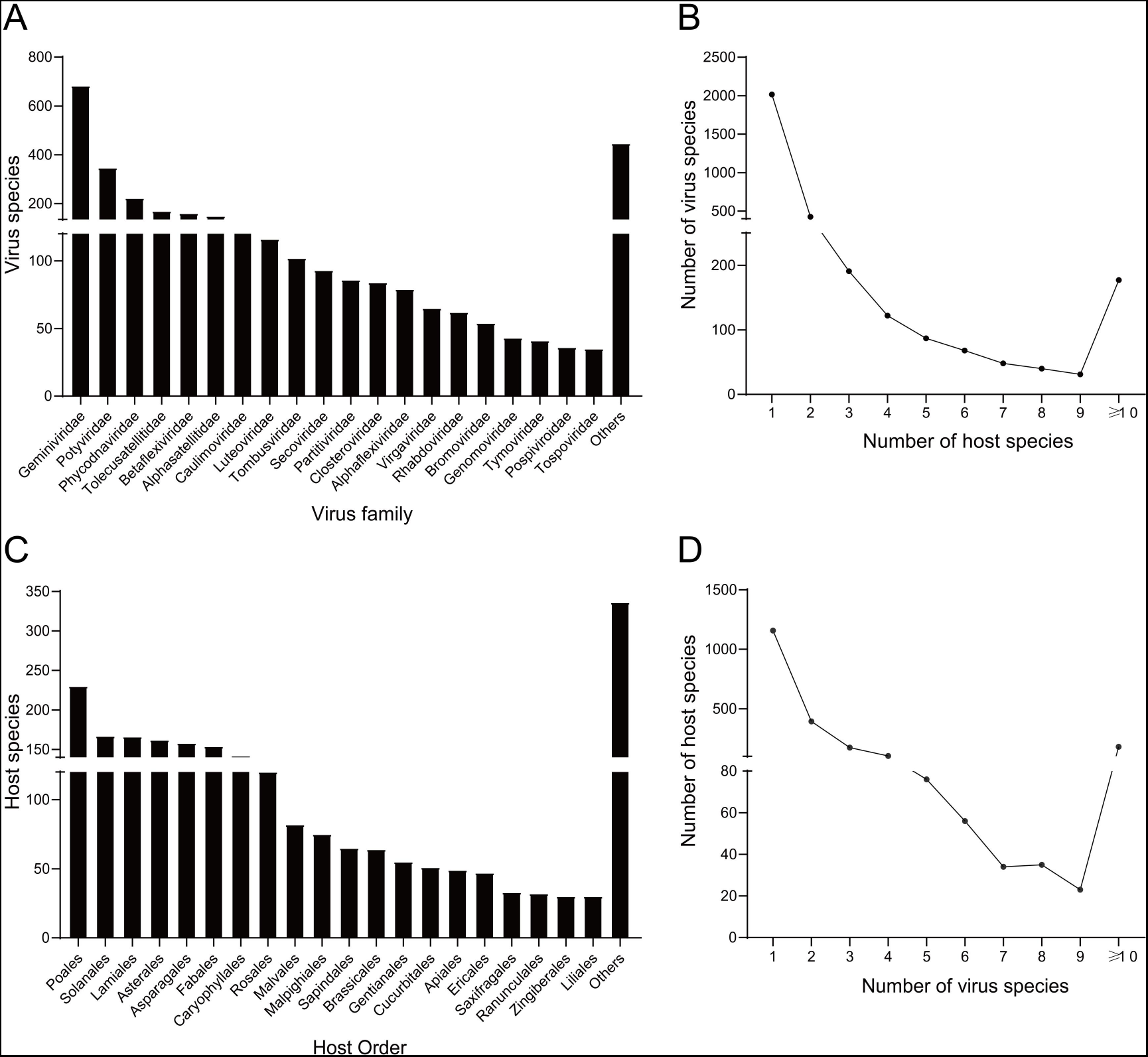
Summary of the viruses, plants and virus-plant interactions used in this study. (A) The number of virus species contained in viral families; (B) The distribution of the number of plant species infected by each virus species; (C) The number of plant species contained in plant orders; (D) The distribution of the number of viral species infecting each plant species.

The 2,231 plant species belonged to 92 plant orders. 97% of plant species belonged to the seed plant, and the remaining plant species belonged to algae, bryophytes and ferns. Most (99%) of seed plants belonged to the angiosperm. The order of Poales contained the largest number of plant species. The top 20 orders contained over 85% of all plant species (Figure 3C). Over 1,000 plant species were infected by only one virus species and 92% of plant species were infected by less than 10 viral species (Figure 3D). Some plant species were infected by more than 10 viral species. For example, the plant species *Solanum lycopersicum* were infected by 419 virus species from 23 viral families.

Analysis of the isolation source of plant viruses by tissue showed that over 92% of virus species were identified from leaves (Figure S3). The remaining virus species were isolated from the stem, root, seed, fruit, flower and other tissues.

### Characterizing the virome in the tropical and temperate plants

Previous studies by geographic and phylogenetic analyses suggest an “out-of-the-tropics” scenario of angiosperm diversification. The ancestral angiosperms prefer warm and humid climates, and the subsequent angiosperms experience transitions into more temperate and arid climates [41]. Therefore, the virome identified in the tropical and temperate plants were analyzed and compared (Figure 4A). A total of 628 and 177 plant species were reported to be only in the tropical and temperate regions, respectively, which were defined as the tropical and temperate plants, respectively. A total of 1,176 virus species from 38 viral families were identified in tropical plants, which were much larger than those identified in temperate plants (281 virus species in 28 viral families). Only 41% virus species identified in the temperate plants were also identified in tropical plants (Figure 4B and Table S1). The composition of viral families identified in the tropical and temperate plants also differed a lot. For example, in the temperate plants, the viral family of *Potyviridae* was most abundant with a ratio of 20.6% which was decreased to 14.5% in the tropical plants; while in the tropical plants, the viral family of *Geminiviridae* was most abundant with a ratio of 22.6% which was decreased to 9.3% in the temperate plants.

**Figure 4.**
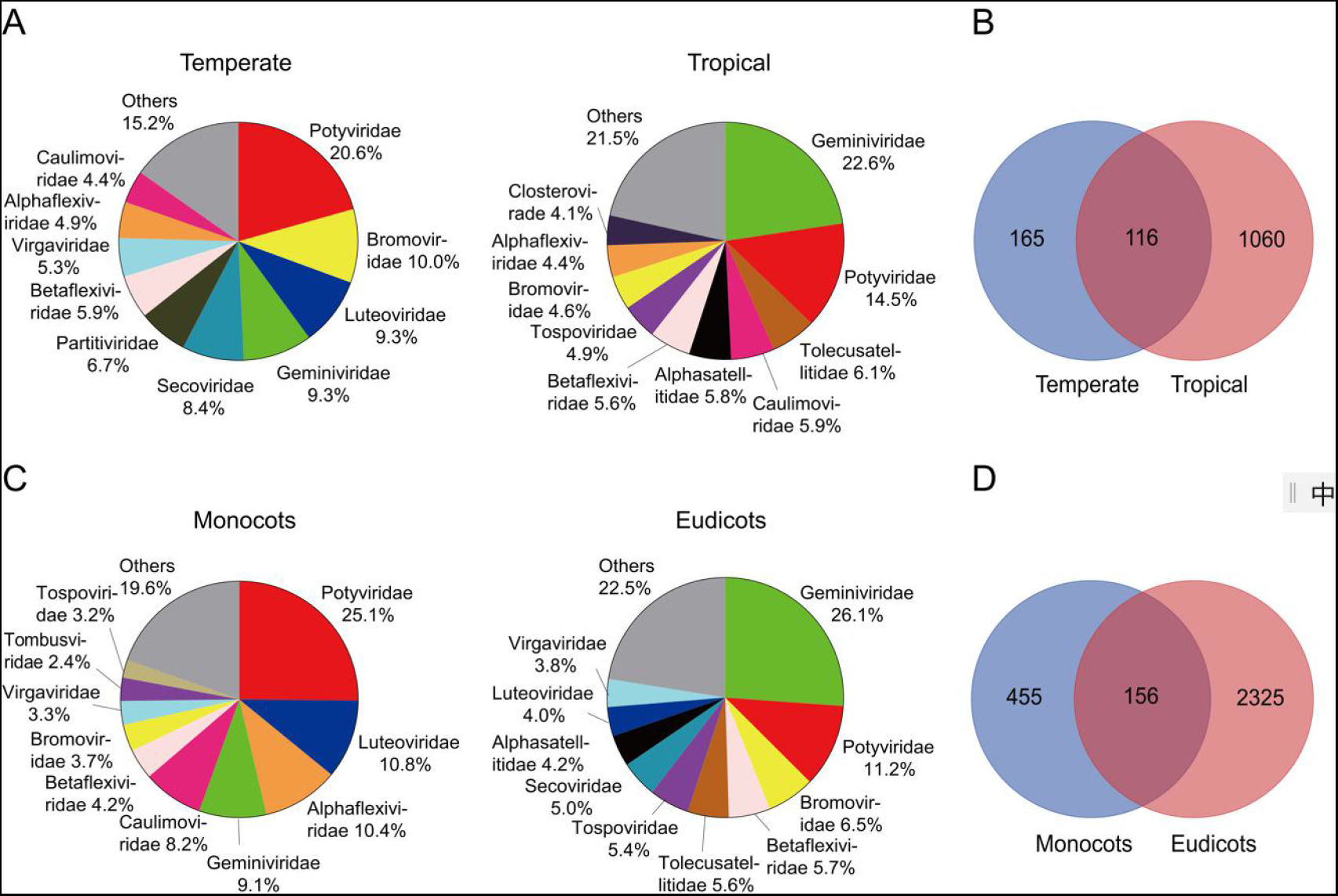
Comparison of the virus family composition between viromes in the Monocots and Eudicots and in the temperate and tropical plants. (A) Composition of viral families identified in the temperate and tropical plants; (B) Venn diagram of the number of viruses identified in the temperate and tropical plants; (C) Composition of viral families identified in the Monocots and Eudicots; (D) Venn diagram of the number of viruses identified in the Monocots and Eudicots.

### Evolution of viromes within the angiosperm

Since most of virus species used in the study were identified in the angiosperm, we focused on the evolution of virome in the angiosperm. There were a total of 49 orders within the angiosperm, including 9 orders in Monocots, 34 orders in Eudicots and 6 orders in early angiosperms (Figure S4). The ratios of the most abundant ten viral families identified in the angiosperm were shown for each plant order (Figure S4). The virome within the Monocots and Eudicots were analyzed and compared (Figure 4C). There were a total of 611 virus species in 37 viral families identified in Monocots, and 2,481 virus species in 42 viral families identified in Eudicots. Most of viral families identified in the Monocots were also identified in the Eudicots (Table S2), although the composition of viral families identified in the Monocots and Eudicots differed a lot (Figure 4C). For example, in the Monocots, the viral family of *Potyviridae* was most abundant with a ratio of 25.1% which was decreased to 11.2% in the Eudicots; while in the Eudicots, the viral family of *Geminiviridae* was most abundant with a ratio of 26.1% which was decreased to 9.1% in the Monocots. In terms of viral species, only 26% of viruses identified in the Monocots were also identified in the Eudicots (Figure 4D).

### Factors associated with the host range of plant viruses

Plant viruses infected a wide range of plant species, albeit most virus species infected fewer than ten plant species. Previous studies have shown that both the genetic and ecological factors contribute to the host range of plant viruses. This study analyzed several factors which may influence the host range of plant viruses and were listed as follows: 1) virus group. The plant viruses used in this study included 1,465 DNA and 1,739 RNA virus species. The RNA viruses were found to have a similar host range to that of DNA viruses (p-value > 0.05 in the Wilcoxon rank-sum test) (Figure 5A); 2) enveloped or not. The plant viruses used in this study included 309 enveloped and 2,175 non-enveloped virus species. The non-enveloped viruses were found to have a slightly larger host range than that of enveloped viruses (p-value < 0.05 in the Wilcoxon rank-sum test) (Figure 5B); 3) genome size and GC content. No significant correlations were observed between the genome size or GC content of viral genomes and the host range (Figure S5); 4) Transmission mode. Plant viruses are transmitted mainly by horizontal transmission and (or) vertical transmission [42]. The viruses were divided into three groups including the horizontal and vertical transmission groups, and the group with both transmission modes. No significant differences were observed between the host range of three groups of viruses (Figure 5C).

**Figure 5.**
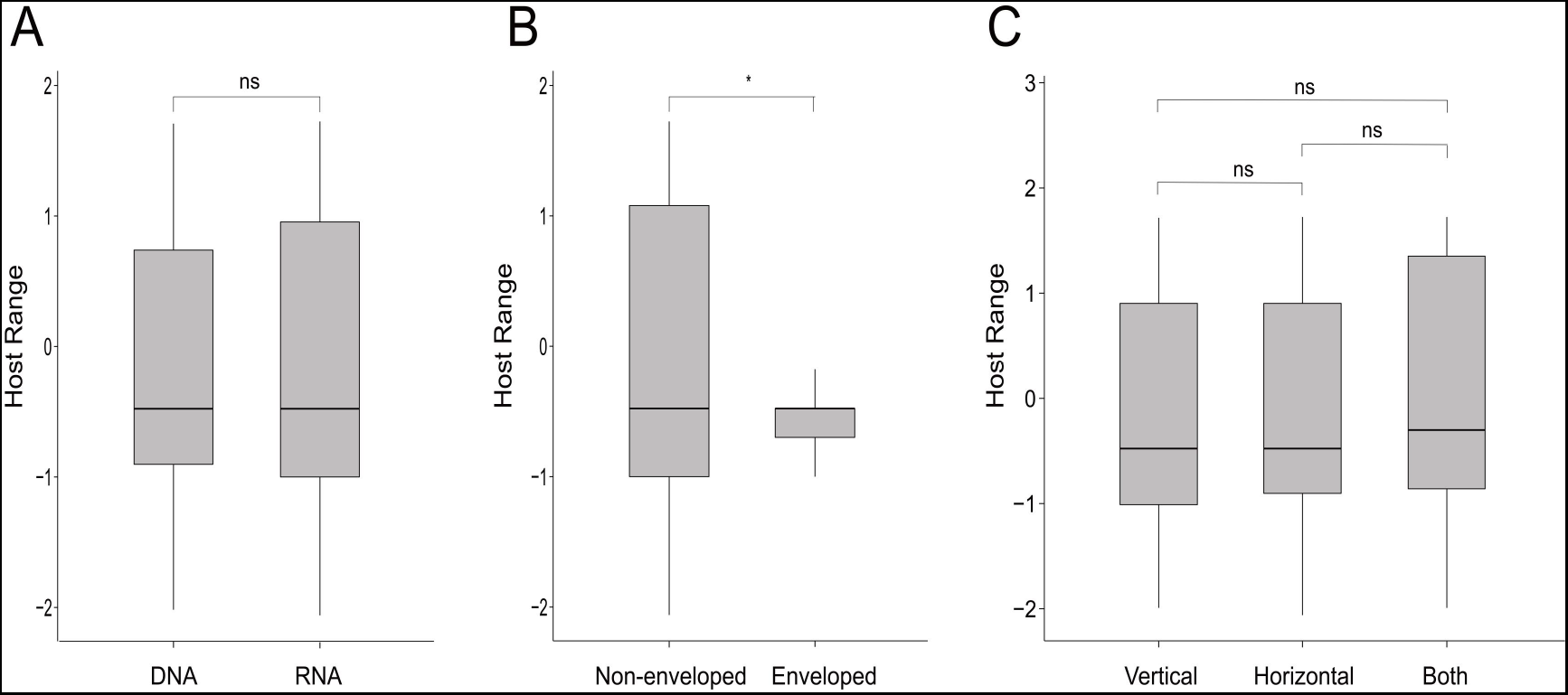
Factors associated with the host range of plant viruses. (A)-(C) refers to the factors of virus group, enveloped or not and transmission mode. For clarity, the host range was transformed with the natural logarithm. *, p-value < 0.05; ns, no significant difference.

### Overview of the Plant Virus Database

A database named Plant Virus Database (PVD) was created to store and organize the plant viruses and it was freely available at http://47.90.94.155/PlantVirusBase/#/home. The PVD mainly includes Home, Browse, Statistic, Download and Help pages (Figure 6).

**Figure 6.**
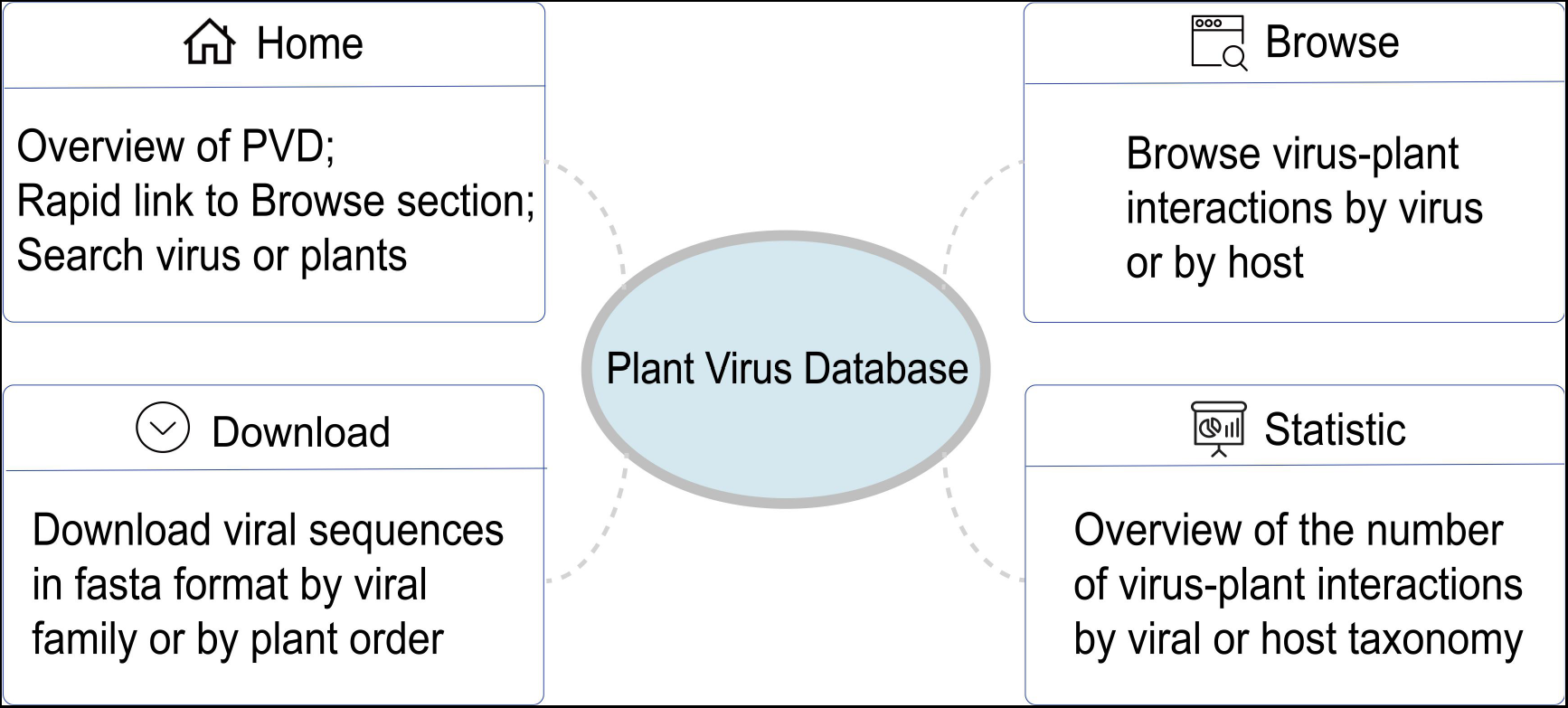
The structure of the Plant Virus Database.

#### Home

The home page provides an introduction of the Plant Virus Database and rapid links to the *Browse* page by plant or viral family. Besides, a search box was also provided for searching virus or plant in the database.

#### Browse

The page displayed the interactions between plant viruses and their hosts. When browsing by viruses, a table was provided which showed the name, taxonomy and hosts of viruses. The viruses shown in the table could be easily filtered by virus name, taxonomy and hosts. When browsing by hosts, a table was provided which showed the name, taxonomy and viruses of the host. The hosts shown in the table could be easily filtered by host name, taxonomy and viruses which infect the host.

#### Statistic

This page displayed a summary statistics about the number of virus-host interactions in each virus or plant taxonomic unit.

#### Download

All the virus genomic sequences were available for downloading by viral family and by plant order.

## Discussion

The study systematically identified and curated an up-to-date atlas of plant viruses and further built the Plant Virus Database (PVD) for storing these viruses and their interactions with plant hosts. Compared to existing plant virus databases such as DPVweb, PVD provided a much larger number of plant virus species (3,206 *vs* 1,503) and virus-host interactions (9,604 *vs* 3,706), and should be a valuable resource for exploring the diversity of plant viruses.

Most plant viruses were observed to only infect a few plants and most plants were also observed to be infected by only a few viruses. Besides, no or few plant viruses were identified in most RNA-seq datasets in the 1KP project, while lots of animal viruses were identified in the RNA-seq datasets from animal samples [20,21]. This may be caused by several factors. Firstly, the plant viruses and virus-plant interactions were less-studied compared to those of animal viruses [43], which results in the under-estimated plant viruses in public databases such as the NCBI RefSeq database on which the workflow developed here were based; secondly, the ability of the workflow used in the study in identifying novel plant viruses is limited as no novel plant viruses were identified in the 1KP project; thirdly, the plants have strong defenses against viral infections and the cell wall of plants can limit access of viral entry into plant cells [44], which may lead to less prevalence of plant viruses than the animal viruses. Nevertheless, the RNA-seq technology is a powerful tool for discovering novel viruses as was demonstrated in numerous studies [19,45]. Our analysis also showed that most of the virus-plant interactions identified in the 1KP project were novel compared to those in public databases (Figure 2), suggesting that the workflow developed here should be effective in identifying viruses from the RNA-seq data although its ability to identify novel viruses needs improvement. Much more viruses and virus-plant interactions would be identified by the workflow giving more RNA-seq datasets are provided.

The role of the virus in plants and plant evolution is unknown yet [13]. Virus infection of plants and host defenses reciprocally affect both the virus and host. To fight against viral infection, the plants have developed diverse strategies [15] such as resistance genes [46]. During the process, the plants also change their phenotypes [47]. This leads to the co-evolution between virus and plants [13]. Therefore, we compared the viromes in the Monocots and Eudicots, and those in the tropical and temperate plants, and found large differences between them. This suggests that viruses may play an important role in the diversification of plants.

Many factors were reported to affect the plant virus host range [16,48]. Firstly, the genetic factors are the main determinants of the virus host range. Genetic specificity is commonly observed in plant viruses [16,49]. That’s to say, only some viruses can infect and multiply in some hosts. Our study observed similar host range between RNA and DNA viruses, and no associations between the length or GC content of viral genomes and the host range, although the non-enveloped viruses were observed to have a slightly larger host range than the enveloped viruses. This suggests that the genetic factors determining the host range of plant viruses need to be determined in the future studies. Secondly, environmental and ecological factors also influence the plant virus host range [16]. For example, the structure of the host species population may have a significant influence on the plant virus host range [18]. When host species co-exist in the same community, and (or) are closely related, the opportunity [50] for viruses to interact with them is more likely than between unrelated hosts in other communities [16,51]. Besides, the stochastic fluctuations in community composition [48], local extinctions and speciation, and the transmission mode of viruses may also affect the virus host range [16]. However, we found that viruses with different transmission mode have no significant difference in host range. The role of the transmission mode in determining the host range of plant viruses need further studies.

There are some limitations in this study. Firstly, the virus-plant interactions identified in the 1KP project are likely to be biased. The absence of virus in the RNA-seq dataset from a certain plant species does not necessarily indicate that the plant will not be infected by this virus. Viruses may infect only certain specific tissues such as the root or the seed which may be not covered in the 1KP project; secondly, the number of plant viruses identified in the study is still limited in its size due to the limited data in the public databases. More efforts are needed to identify plant viruses; thirdly, the workflow developed here needs improvement in its ability to identify novel viruses; fourthly, most plant viruses were identified based on the sequencing technology. More experimental efforts are needed to validate these plant viruses. Nevertheless, this study provided an up-to-date resource of plant viruses. It has the potential to improve our knowledge of the plant virus genetic diversity and the virus-host interactions.

## Supporting information

Supplementary materals

## Key Points

- More than 600 pairs of virus-plant interactions were identified from the 1KP project.
- The PVD contained 3,206 virus species and 9,604 virus-plant host interactions which were more than twice that reported in previous plant virus databases.
- Significant different virome compositions were observed between Monocots and Eudicots, and between plants in tropical and temperate regions.

## ACKNOWLEDGEMENTS

This work was supported by the Hunan Provincial Natural Science Foundation of China (2020JJ3006) and National Natural Science Foundation of China (32170651).

## Compliance with Ethical Standards

### Animal and Human Rights Statement

This article does not contain any studies with human or animal subjects performed by the author.

## References

1. Zaitlin M, Palukaitis P. Advances in Understanding Plant Viruses and Virus Diseases. Annu Rev Phytopathol 2000;38:117–143.

2. Hull R. Chapter 4 - Symptoms and Host Range In: Hull R, ed. Plant Virology (Fifth Edition). Boston: Academic Press, 2014;145–198.

3. Jones RA. Control of plant virus diseases. Adv Virus Res 2006;67:205–244.

4. Mohammed Riyaz SU, Immanuel Jesse DM, Kathiravan K. Chapter 12 - Plant virus: Diversity and ecology In: Gaur RK, Khurana SMP, Sharma P, et al., eds. Plant Virus-Host Interaction (Second Edition). Boston: Academic Press, 2021;319–328.

5. Shafiq M, Qurashi F, Mushtaq S, et al. Chapter 13 - DNA plant viruses: biochemistry, replication, and molecular genetics In: Awasthi LP, ed. Applied Plant Virology: Academic Press, 2020;169–182.

6. Gergerich RC, Dolja VV. Introduction to Plant Viruses, the Invisible Foe. Plant Health Instructor 2006.

7. Lucas WJ. Plant viral movement proteins: Agents for cell-to-cell trafficking of viral genomes. Virology 2006;344:169–184.

8. Mauck KE, Chesnais Q, Shapiro LR. Evolutionary Determinants of Host and Vector Manipulation by Plant Viruses. Adv Virus Res 2018;101:189–250.

9. Belshaw R, Gardner A, Rambaut A, et al. Pacing a small cage: mutation and RNA viruses. Trends in Ecology & Evolution 2008;23:188–193.

10. Hadidi A, Flores R, Randles JW, et al. Viroids and Satellites. Viroids and Satellites. Boston: Academic Press, 2017.

11. Diener TO. Viroids: the smallest known agents of infectious disease. Annu Rev Microbiol 1974;28:23–39.

12. Francki RI. Plant virus satellites. Annu Rev Microbiol 1985;39:151–174.

13. Fraile A, García-Arenal F. Chapter 1 - The Coevolution of Plants and Viruses: Resistance and Pathogenicity In: Carr JP, Loebenstein G, eds. Advances in Virus Research: Academic Press, 2010;1–32.

14. Garcia-Ruiz H. Host factors against plant viruses. Mol Plant Pathol 2019;20:1588–1601.

15. Avinash M, Rajarshi Kumar G. Host Plant Strategies to Combat Against Viruses Effector Proteins. Current Genomics 2020;21:401–410.

16. McLeish MJ, Fraile A, García-Arenal F. Evolution of plant-virus interactions: host range and virus emergence. Curr Opin Virol 2019;34:50–55.

17. Monjane AL, Dellicour S, Hartnady P, et al. Symptom evolution following the emergence of maize streak virus. eLife 2020;9:e51984.

18. Elena SF, Fraile A, García-Arenal F. Chapter Three - Evolution and Emergence of Plant Viruses In: Maramorosch K, Murphy FA, eds. Advances in Virus Research: Academic Press, 2014;161–191.

19. Cantalupo PG, Pipas JM. Detecting viral sequences in NGS data. Current Opinion in Virology 2019;39:41–48.

20. Shi M, Lin X-D, Tian J-H, et al. Redefining the invertebrate RNA virosphere. Nature 2016;540:539–543.

21. Shi M, Lin XD, Chen X, et al. The evolutionary history of vertebrate RNA viruses. Nature 2018;556:197–202.

22. Lee HK, Kim SY, Yang HJ, et al. The Detection of Plant Viruses in Korean Ginseng (Panax ginseng) through RNA Sequencing. Plant Pathol J 2020;36:643–650.

23. Rumbou A, Candresse T, Marais A, et al. Unravelling the virome in birch: RNA-Seq reveals a complex of known and novel viruses. PLoS One 2020;15:e0221834.

24. Carpenter EJ, Matasci N, Ayyampalayam S, et al. Access to RNA-sequencing data from 1,173 plant species: The 1000 Plant transcriptomes initiative (1KP). GigaScience 2019;8.

25. Adams MJ, Antoniw JF. DPVweb: a comprehensive database of plant and fungal virus genes and genomes. Nucleic Acids Res 2006;34:D382–385.

26. Brunt AA, Crabtree K, Dallwitz MJ, et al. Plant Viruses Online: Descriptions and Lists from the VIDE Database. Online Publication. 1996.

27. Roenhorst JW, Boonham N, Winter S, et al. The plant viruses and viroids database and collections of Q-bank. EPPO Bulletin 2013;43:238–243.

28. Sayers EW, Cavanaugh M, Clark K, et al. GenBank. Nucleic Acids Res 2021;49:D92–d96.

29. Mihara T, Nishimura Y, Shimizu Y, et al. Linking Virus Genomes with Host Taxonomy. Viruses 2016;8:66.

30. Chen S, Zhou Y, Chen Y, et al. fastp: an ultra-fast all-in-one FASTQ preprocessor. Bioinformatics 2018;34:i884–i890.

31. Grabherr MG, Haas BJ, Yassour M, et al. Full-length transcriptome assembly from RNA-Seq data without a reference genome. Nat Biotechnol 2011;29:644–652.

32. Li B, Dewey CN. RSEM: accurate transcript quantification from RNA-Seq data with or without a reference genome. BMC Bioinformatics 2011;12:323.

33. O’Leary NA, Wright MW, Brister JR, et al. Reference sequence (RefSeq) database at NCBI: current status, taxonomic expansion, and functional annotation. Nucleic Acids Res 2016;44:D733–745.

34. Altschul SF, Gish W, Miller W, et al. Basic local alignment search tool. J Mol Biol 1990;215:403–410.

35. Marchler-Bauer A, Anderson JB, Derbyshire MK, et al. CDD: a conserved domain database for interactive domain family analysis. Nucleic Acids Res 2007;35:D237–240.

36. Sayers EW, Beck J, Bolton EE, et al. Database resources of the National Center for Biotechnology Information. Nucleic Acids Res 2021;49:D10–d17.

37. Shen W, Ren H. TaxonKit: A practical and efficient NCBI taxonomy toolkit. J Genet Genomics 2021.

38. Federhen S. The NCBI Taxonomy database. Nucleic Acids Res 2012;40:D136–143.

39. Zy W, Zhou Z-K, Li D-Z, et al. The areal types of the world families of seed plants. Acta Botanica Yunnanica 2003;25:245–257.

40. Group TAP. An update of the Angiosperm Phylogeny Group classification for the orders and families of flowering plants: APG IV. Botanical Journal of the Linnean Society 2016;181:1–20.

41. Ramírez-Barahona S, Sauquet H, Magallón S. The delayed and geographically heterogeneous diversification of flowering plant families. Nature Ecology & Evolution 2020;4:1232–1238.

42. Stevens WA. Transmission of Plant Viruses. Virology of Flowering Plants. Boston, MA: Springer US, 1983;41–68.

43. Ingwell LL, Eigenbrode SD, Bosque-Pérez NA. Plant viruses alter insect behavior to enhance their spread. Sci Rep 2012;2:578.

44. Buchmann JP, Holmes EC. Cell Walls and the Convergent Evolution of the Viral Envelope. Microbiology and Molecular Biology Reviews 2015;79:403–418.

45. Maliogka VI, Minafra A, Saldarelli P, et al. Recent Advances on Detection and Characterization of Fruit Tree Viruses Using High-Throughput Sequencing Technologies. Viruses 2018;10.

46. Ishibashi K, Ishikawa M. [Virus resistance genes in plants]. Uirusu 2018;68:13–20.

47. Mauck KE. Variation in virus effects on host plant phenotypes and insect vector behavior: what can it teach us about virus evolution? Current Opinion in Virology 2016;21:114–123.

48. McLeish MJ, Fraile A, García-Arenal F. Chapter Nine - Ecological Complexity in Plant Virus Host Range Evolution In: Malmstrom CM, ed. Advances in Virus Research: Academic Press, 2018;293–339.

49. Vassilakos N, Simon V, Tzima A, et al. Genetic Determinism and Evolutionary Reconstruction of a Host Jump in a Plant Virus. Molecular Biology and Evolution 2015;33:541–553.

50. Kedem H, Cohen C, Messika I, et al. Multiple effects of host-species diversity on coexisting host-specific and host-opportunistic microbes. Ecology 2014;95:1173–1183.

51. Moury B, Fabre F, Hébrard E, et al. Determinants of host species range in plant viruses. J Gen Virol 2017;98:862–873.

