## Supplementary materals for "Plant Virus Database: a resource for exploring the diversity of plant viruses and their interactions with hosts"

### Supplementary Figures

**Figure S1** Distribution of the number of viral species identified per RNA-seq dataset in the 1KP project.

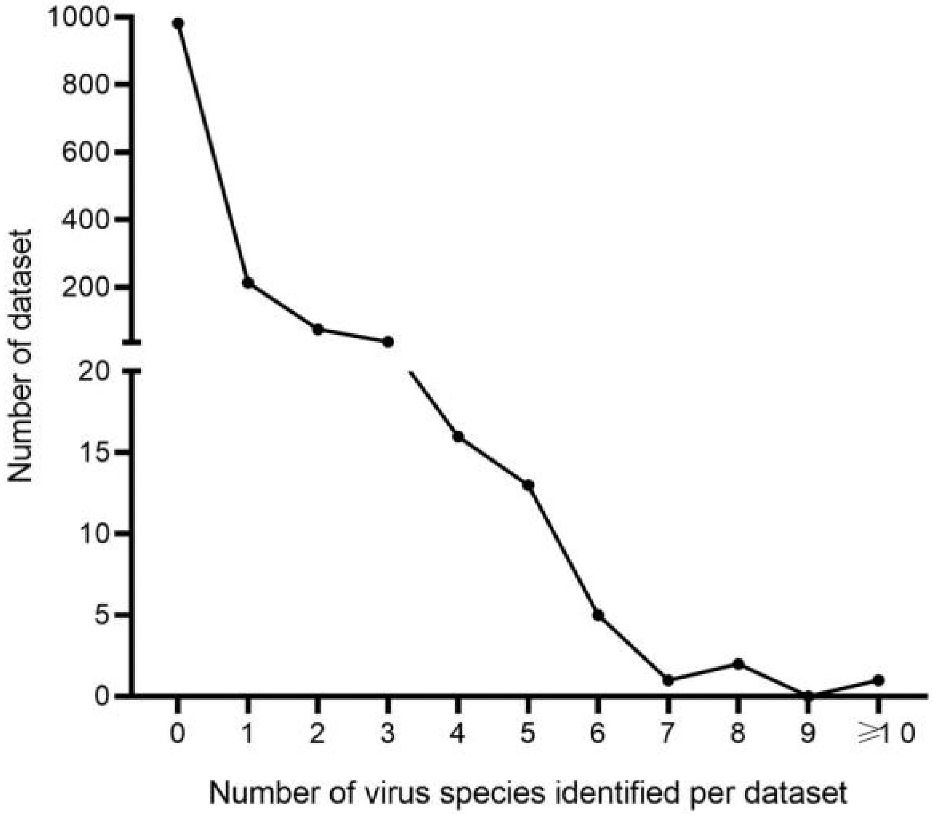

**Figure S2** Summary of the viruses, plants and virus-plant interactions obtained from the 1KP project. (A) The number of viruses identified in viral families; (B) The distribution of the number of plant species infected by viruses; (C) The number of plant species in plant orders; (D) The distribution of the number of viral species infecting each plant species.

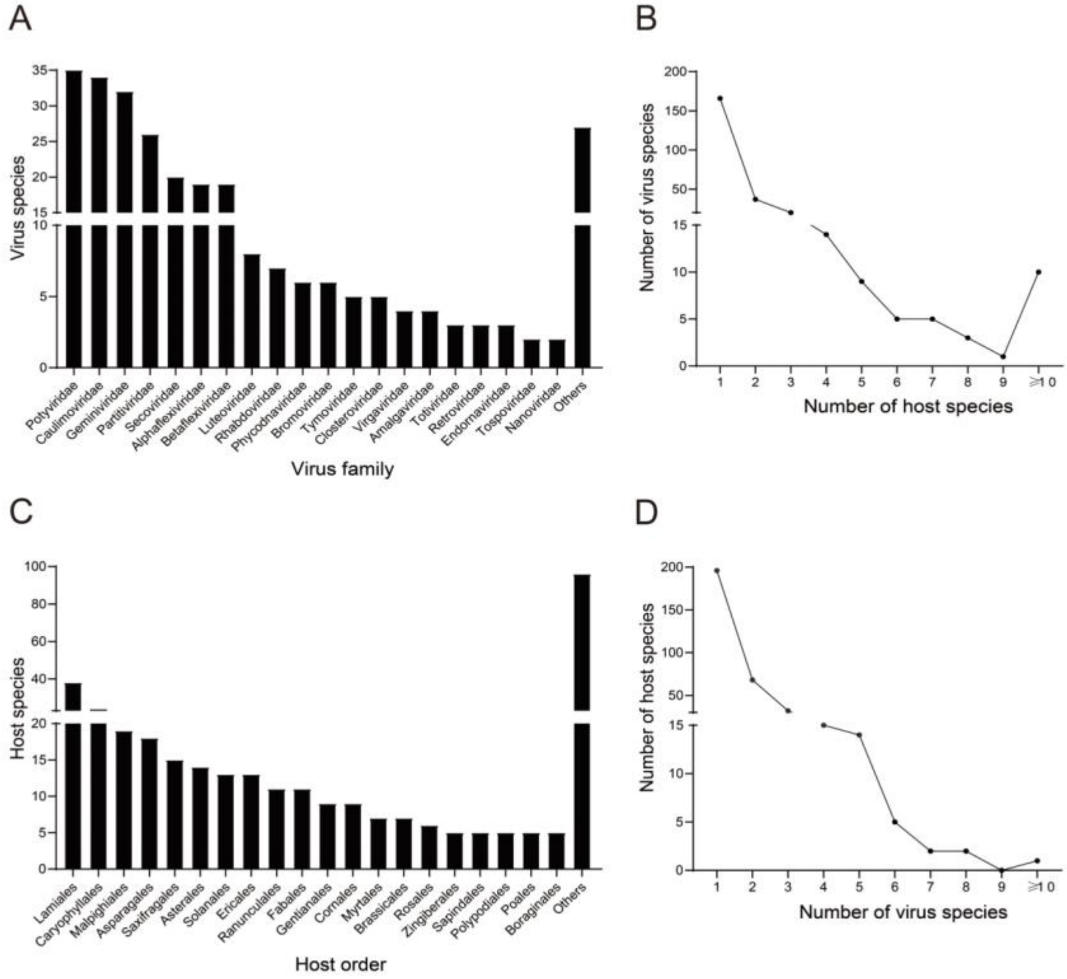

**Figure S3** Isolation source of plant viruses by tissue.

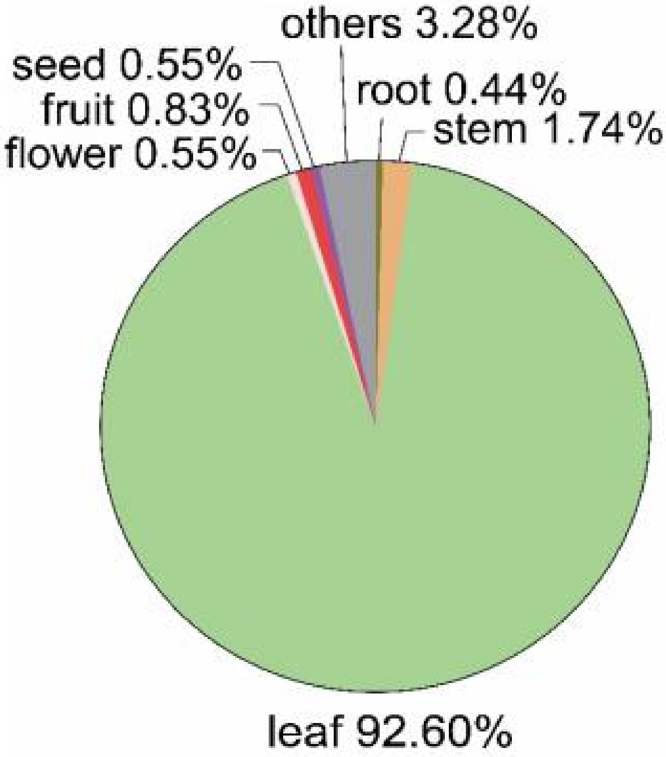

**Figure S4** The ratios of the most abundant ten viral families identified in the angiosperm in 49 plant orders. The viral family was colored according to the legend in the bottom-right. The number besides the bar showed the number of virus species identified in the order. The asterisks referred to the plant orders which were mainly located in the temperate regions.

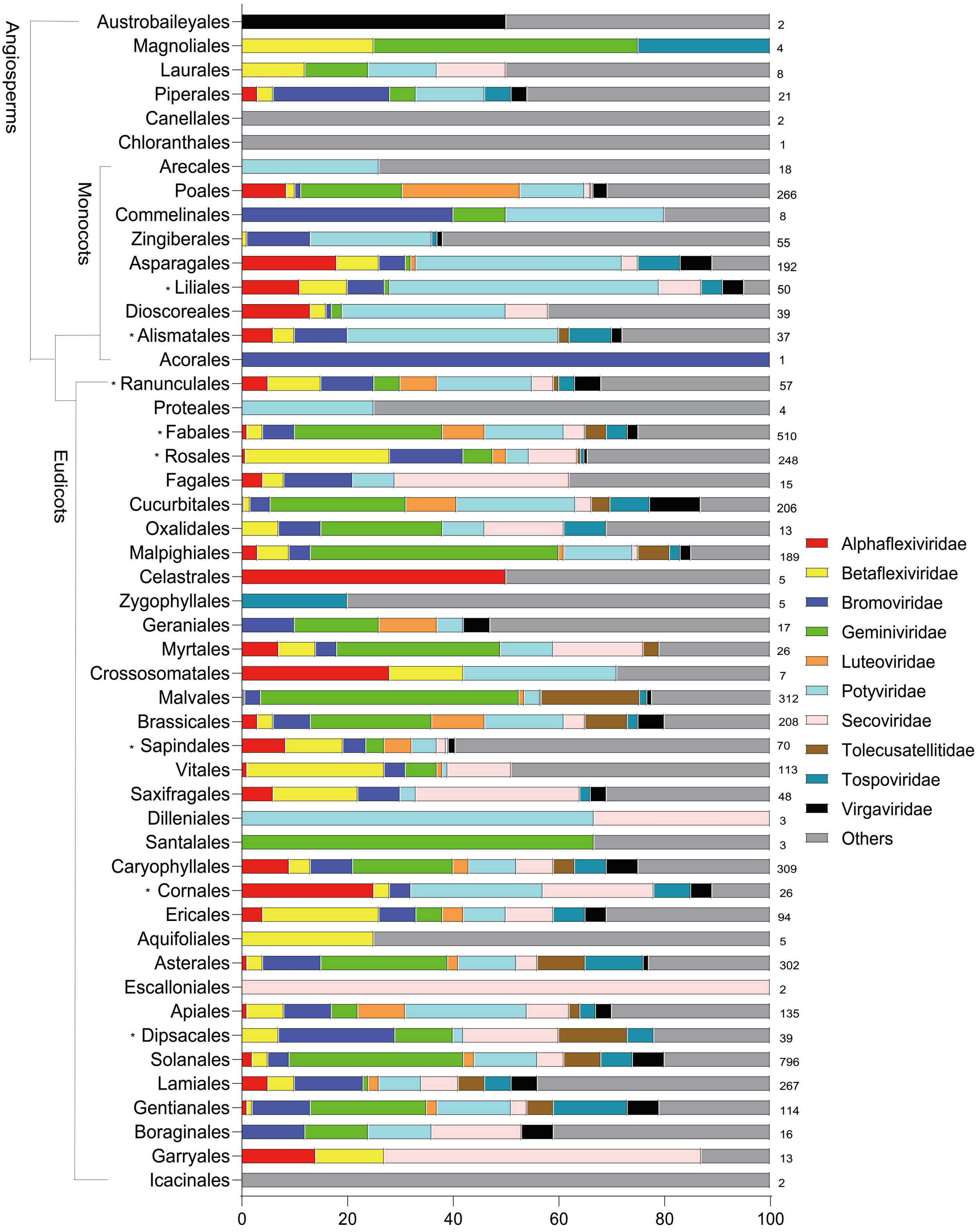

**Figure S5** The relationship between the host range and the viral genome length (A) and the GC content of viral genomes (B). The Pearson correlation coefficients (PCC) and the related p-values were shown in the top-right.

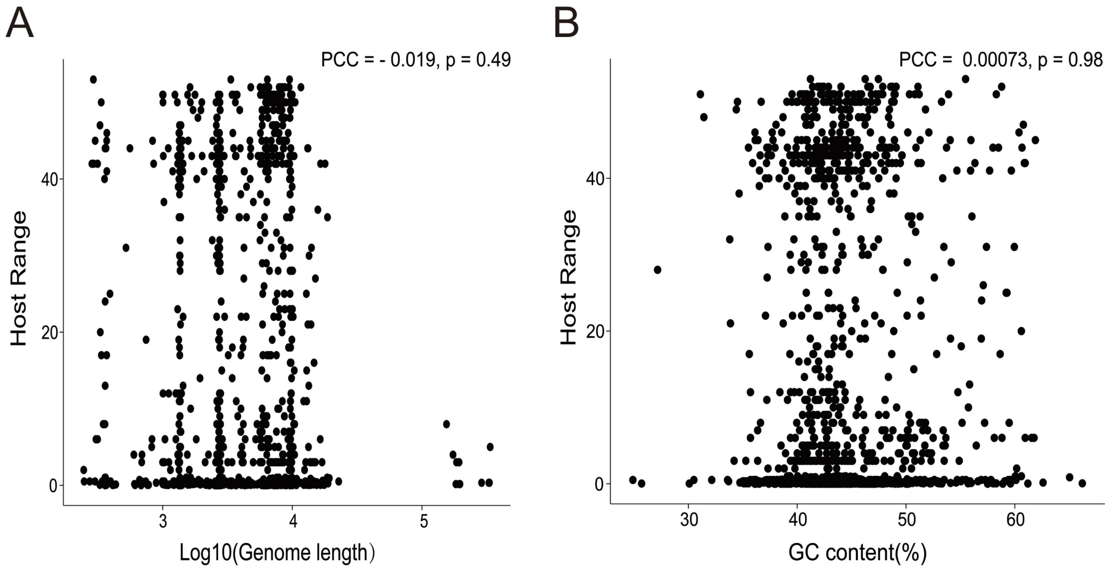


### Supplementary Tables

**Table S1** The host range of viruses in the viral families which infect the plants in temperate and tropical regions.

| Virus Family | Total number of virus species in the family | Number of virus species infecting temperate plants only | Number of virus species infecting tropical plants only | Number of viruses infecting both temperate and tropical plants |
|---|---|---|---|---|
| Alphaflexiviridae | 61 | 6 | 47 | 8 |
| Alphasatellitidae | 81 | 2 | 73 | 6 |
| Amalgaviridae | 6 | 2 | 4 | 0 |
| Aspiviridae | 2 | 0 | 2 | 0 |
| Avsunviroidae | 2 | 1 | 1 | 0 |
| Benyviridae | 2 | 0 | 2 | 0 |
| Betaflexiviridae | 80 | 14 | 59 | 7 |

| | | | | |
|---|---|---|---|---|
| Botourmiaviridae | 1 | 0 | 1 | 0 |
| Bromoviridae | 25 | 9 | 10 | 6 |
| Caulimoviridae | 85 | 4 | 72 | 9 |
| Chrysoviridae | 2 | 2 | 0 | 0 |
| Closteroviridae | 40 | 1 | 36 | 3 |
| Endornaviridae | 9 | 0 | 9 | 0 |
| Fimoviridae | 10 | 2 | 8 | 0 |
| Gammaflexiviridae | 1 | 0 | 1 | 0 |
| Geminiviridae | 276 | 13 | 239 | 24 |
| Genomoviridae | 17 | 3 | 14 | 0 |
| Hypoviridae | 1 | 0 | 1 | 0 |
| Kitaviridae | 6 | 0 | 6 | 0 |
| Luteoviridae | 29 | 7 | 19 | 3 |
| Mayoviridae | 2 | 1 | 1 | 0 |
| Mitoviridae | 3 | 0 | 1 | 0 |
| Nanoviridae | 7 | 1 | 5 | 1 |
| Partitiviridae | 45 | 22 | 19 | 4 |
| Parvoviridae | 3 | 0 | 3 | 0 |
| Phenuiviridae | 9 | 1 | 8 | 0 |
| Pospiviroidae | 22 | 1 | 18 | 3 |
| Potyviridae | 185 | 28 | 141 | 16 |
| Reoviridae | 1 | 0 | 1 | 0 |
| Retroviridae | 3 | 0 | 3 | 0 |
| Rhabdoviridae | 22 | 1 | 20 | 1 |
| Secoviridae | 41 | 8 | 22 | 11 |
| Solemoviridae | 4 | 1 | 3 | 0 |
| Tolecusatellitidae | 80 | 7 | 67 | 6 |
| Tombusviridae | 34 | 10 | 22 | 2 |
| Tospoviridae | 16 | 0 | 14 | 2 |
| Totiviridae | 4 | 0 | 4 | 0 |
| Tymoviridae | 16 | 3 | 13 | 0 |
| Virgaviridae | 32 | 4 | 22 | 6 |

**Table S2** The host range of viruses in the viral families which infect the Monocots and Eudicots.

| Virus Family | Total number of virus species in the family | Number of virus species infecting Monocots only | Number of virus species infecting Eudicots only | Number of viruses infecting both Monocots and |
|---|---|---|---|---|

| | | | | Eudicots |
|---|---|---|---|---|
| Alphaflexiviridae | 79 | 25 | 34 | 20 |
| Alphasatellitidae | 145 | 12 | 125 | 8 |
| Amalgaviridae | 24 | 12 | 12 | 0 |
| Aspiviridae | 7 | 0 | 7 | 0 |
| Avsunviroidae | 4 | 0 | 4 | 0 |
| Benyviridae | 6 | 2 | 4 | 0 |
| Betaflexiviridae | 159 | 22 | 131 | 6 |
| Botourmiaviridae | 8 | 1 | 7 | 0 |
| Bromoviridae | 54 | 1 | 47 | 6 |
| Caulimoviridae | 132 | 39 | 82 | 11 |
| Chrysoviridae | 3 | 1 | 2 | 0 |
| Closteroviridae | 84 | 17 | 67 | 0 |
| Deltaflexiviridae | 1 | 1 | 0 | 0 |
| Dicistroviridae | 1 | 0 | 1 | 0 |
| Endornaviridae | 26 | 4 | 21 | 1 |
| Fimoviridae | 27 | 3 | 24 | 0 |
| Gammaflexiviridae | 1 | 0 | 1 | 0 |
| Geminiviridae | 676 | 35 | 635 | 6 |
| Genomoviridae | 43 | 6 | 33 | 4 |
| Hypoviridae | 1 | 0 | 1 | 0 |
| Kitaviridae | 9 | 0 | 8 | 1 |
| Luteoviridae | 116 | 23 | 90 | 3 |
| Mayoviridae | 5 | 1 | 4 | 0 |
| Mitoviridae | 16 | 1 | 15 | 0 |
| Nanoviridae | 16 | 4 | 11 | 1 |
| Nodaviridae | 3 | 1 | 2 | 0 |
| Partitiviridae | 81 | 12 | 66 | 3 |
| Parvoviridae | 5 | 0 | 5 | 0 |
| Peribunyaviridae | 2 | 1 | 1 | 0 |
| Phenuiviridae | 20 | 10 | 10 | 0 |
| Pospiviroidae | 36 | 2 | 33 | 1 |
| Potyviridae | 345 | 117 | 194 | 34 |
| Reoviridae | 13 | 10 | 3 | 0 |
| Retroviridae | 3 | 0 | 2 | 1 |
| Rhabdoviridae | 61 | 15 | 44 | 2 |
| Secoviridae | 93 | 4 | 78 | 11 |
| Solemoviridae | 25 | 7 | 18 | 0 |
| Tolecusatellitidae | 166 | 0 | 164 | 2 |
| Tombusviridae | 101 | 17 | 77 | 7 |
| Tospoviridae | 35 | 0 | 25 | 10 |
| Totiviridae | 12 | 7 | 4 | 1 |
| Tymoviridae | 41 | 4 | 37 | 0 |

| | | | | |
|---|---|---|---|---|
| Virgaviridae | 65 | 7 | 49 | 9 |
